# Nonlinear diversification rates of linguistic phylogenies over the Holocene

**DOI:** 10.1101/553966

**Authors:** Marcus J Hamilton, Robert S. Walker

## Abstract

The expansion of the human species out of Africa in the Pleistocene, and the subsequent development of agriculture in the Holocene resulted in waves of linguistic diversification and replacement across the planet. Analogous to the growth of populations or the speciation of biological organisms, languages diversify over time to form phylogenies of language families. However, the dynamics of this diversification process are unclear. Bayesian methods applied to lexical and phonetic data have created dated linguistic phylogenies for 18 language families encompassing ∼3,000 of the world’s ∼7,000 extant languages. In this paper we use these phylogenies to quantify how fast languages expand and diversify through time both within and across language families. The overall diversification rate of languages in our sample is ∼0.001 yr^-1^ (or a doubling time of ∼700 yr) over the last 6,000 years with evidence for nonlinear dynamics in language diversification rates over time, where both within and across language families, diversity initially increases rapidly and then slows. The large-scale replacement of the world’s hunter-gatherer languages by agricultural languages over the Holocene was a non-constant process.

## Introduction

Following the expansion of modern humans out of Africa ∼60,000 yrs BP, populations and their cultures diversified and there are now over 7,000 languages around the planet [1,2], more than the total number of mammal species [3]. Moreover, most of the world’s current languages belong to agricultural language families [4], and so the majority of current ethnolinguistic diversity has most likely evolved only in the last few thousand years since the Neolithic [5-7]. Somewhat uncertain, however, is the rate at which this diversity evolved [8-11]. Similar questions are asked in the study of biodiversity [12]. Under one scenario, rates of linguistic diversification may slow through time as populations increasingly compete for finite amounts of energy and space in environments, in which case diversification eventually slows and asymptotes toward a quasi-equilibrium. Alternatively, ethnolinguistic diversity may be out of equilibrium because of the relatively recent agricultural revolution, and associated population growth that replaced hunter-gatherer ethnolinguistic diversity over large parts of the planet. Or, perhaps no stable equilibrium exists because between-group competition, or other mechanisms, continuously drive increasing ethnolinguistic diversification and replacement independent of larger environmental constraints (i.e., diversity begets diversity).

To understand how linguistic diversification has played out over time we focus on the structure of linguistic phylogenies. Phylogenetic trees based on linguistic variation have proven to be a powerful analytical tool for reconstructing human population and cultural histories [13-15]. By utilizing a Bayesian statistical approach on the systematic codings of linguistic cognates or phonetic data, these methods have generated dated phylogenies for many of the world’s language families, including Austronesian [13], Bantu [14], and Indo-European [15] (Figure 1). These phylogenies allow us to estimate rates of linguistic diversification, show how they vary through time, and compare rates across different language families.

**Figure 1.**
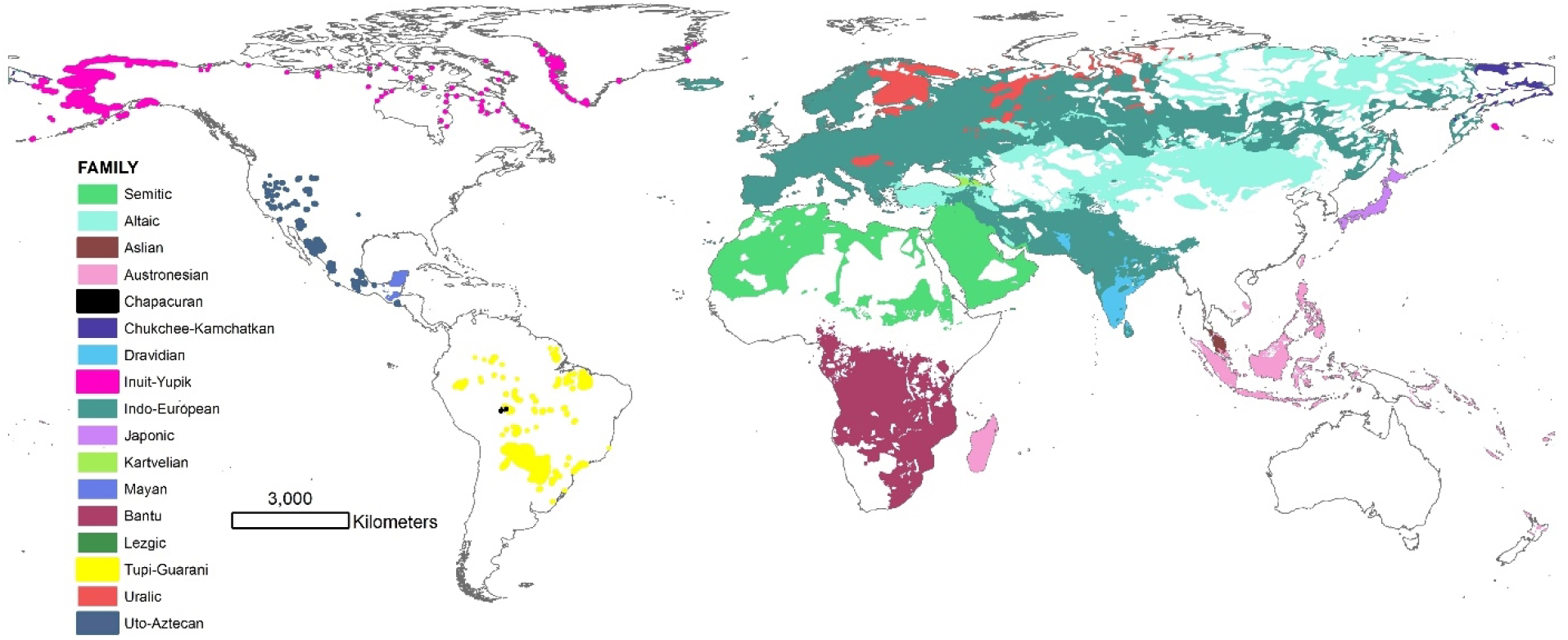
Map of the language families in our sample using polygons from the Ethnologue [2].

In a simple binary branching process, the number of languages, *N*, will increase exponentially with time, *t*, following the exponential growth function, *N* (*t*) = *N*_0_*e*^*rt*^, where *r* = *b* − *d*, the net difference in language speciations, *b* and extinctions, *d*. Starting from the root of a language family, *N*_0_ = 1, and solving for the net diversification rate, *r*, we define

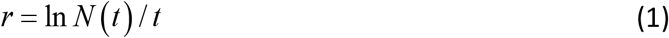

If the slope, *r*, of the number of languages, *N*, through time, *t*, is linear in a semi-log plot then the growth rate is multiplicative and approximately constant, and therefore exponential [16-18] (Figure 2). However, if diversification rates *r* vary through time, either because of time-varying speciation or extinction rates [19, 20], then the slope *r* will not be linear on a semi-log scale, therefore not constant and the growth rate is time-dependent. To explore the growth rates we first plot data for all lineages, and the sum of all lineages, in Figure 2. Second, for each lineage we then examine the distribution of growth rates.

**Figure 2.**
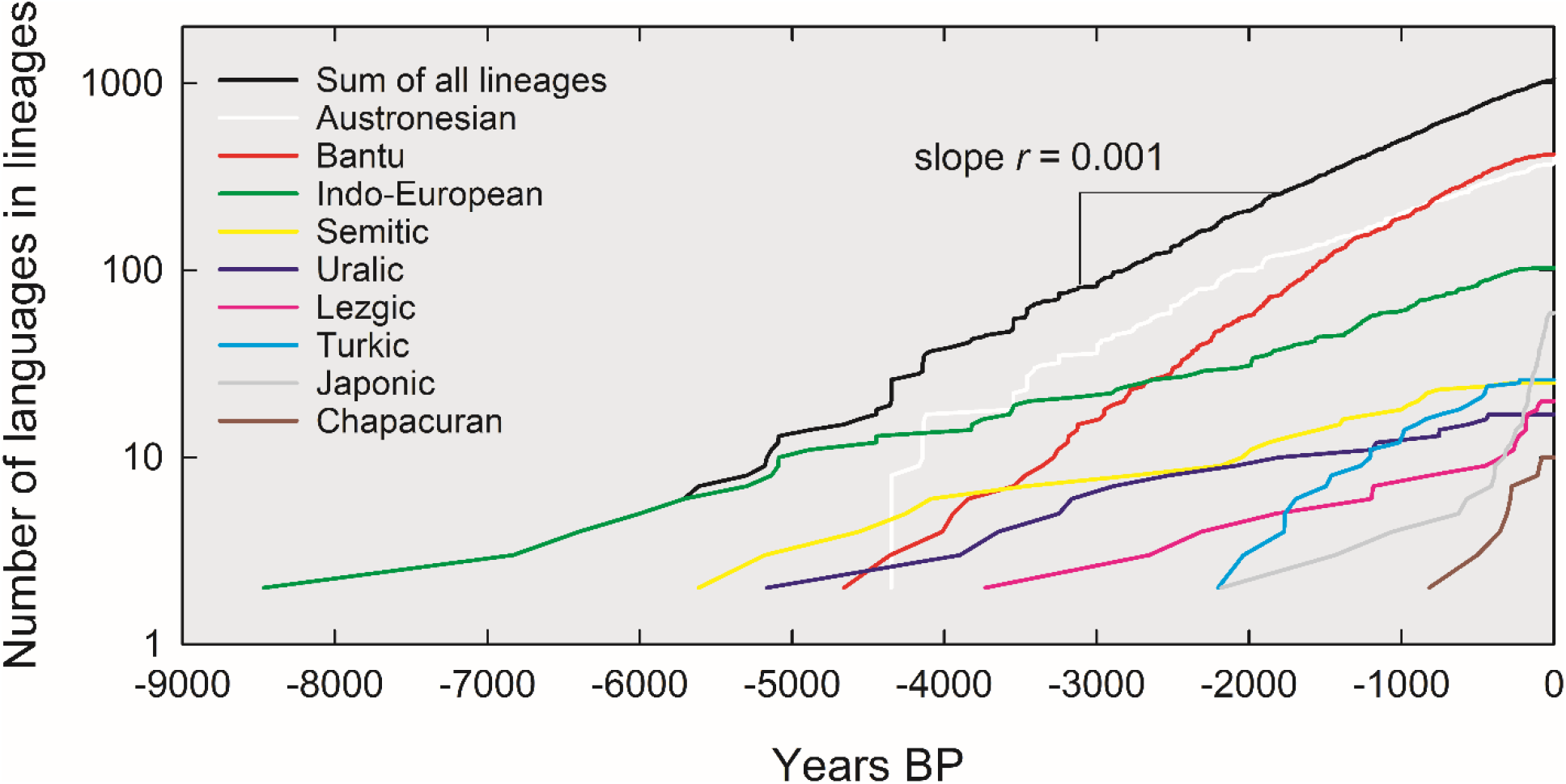
Lineages-through-time plots for 9 language families and the summed total lineages. The number of lineages on the y-axis is on a log scale such that the slopes of these lines are estimates of diversification rate *r*. While there is considerable variation in diversification rates within and across language families, the sum of the lineages suggests an approximately constant diversification rate overall.

The first basic rate of interest for each phylogenetic tree is the overall expansion rate, *r*_*E*_, which is simply the maximum size, *N*_max_ of the *i*th tree divided by its’ time depth, *T*_*i*_ or

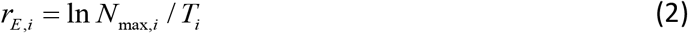

From the overall expansion rate we then establish a doubling time,

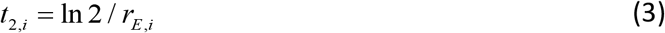

Which is the number of years it takes for the number of languages in the *i*th tree to double (Table 1).

**Table 1.** Language families analyzed in this study sorted by increasing expansion rates, *r*_*E*_. Under data column, “l” indicates languages and “d” dialects. Sample sizes are given for numbers in the available phylogenies, including dialects where applicable, and the total number of extant languages in the family from the Ethnologue [2]. Diversification rates are estimated from equation 1 using only extant languages (natural-logged) divided by estimated root age in years.

| Family/clade | Data | N<sub>sample</sub> | N<sub>extant</sub> | Root age | $r_E$ | Doubling time | Source |
| --- | --- | --- | --- | --- | --- | --- | --- |
| Kartvelian | l |  | 5 | 5500 | 0.00029 | 2369 | 35 |
| Dravidian | l |  | 85 | 13000 | 0.00034 | 2028 | 35 |
| Altaic | l |  | 65 | 11000 | 0.00038 | 1827 | 35 |
| Mayan | l |  | 35 | 6500 | 0.00055 | 1267 | 36 |
| Lezgic | l, d | 20 | 9 | 3730 | 0.00059 | 1177 | 37 |
| Inuit-Yupik | l |  | 11 | 4000 | 0.00060 | 1156 | 35 |
| Uralic | l | 17 | 39 | 5300 | 0.00069 | 1003 | 38 |
| Indo-European | l | 103 | 449 | 8700 | 0.00070 | 987 | 15 |
| Aslian | l, d |  | 18 | 4000 | 0.00072 | 959 | 39 |
| Semitic | l | 25 | 78 | 5614 | 0.00078 | 893 | 40 |
| Uto-Aztecan | l |  | 61 | 5000 | 0.00082 | 843 | 41 |
| Japonic | l, d | 59 | 12 | 2182 | 0.00114 | 609 | 42 |
| Bantu | l | 409 | 668 | 4800 | 0.00136 | 512 | 14 |
| Austronesia | l | 398 | 1268 | 5230 | 0.00137 | 507 | 13 |
| Chapacuran | l, d | 10 | 5 | 1039 | 0.00155 | 447 | 43 |
| Tupi-Guarani | l |  | 51 | 2500 | 0.00157 | 441 | 44 |
| Chukchee-Kamchatkan | l |  | 5 | 1000 | 0.00161 | 431 | 35 |
| Turkic | l | 26 | 41 | 2206 | 0.00168 | 412 | 45 |

In evolutionary biology decreasing rates of species diversification through time are often interpreted as evidence for negative density dependence where ecological niches become saturated limiting opportunities for further speciation [21-26]. Speciation and extinction rates often vary through time leading to expanding or contracting biodiversity and are not well fit by simple exponential growth models [12, 22-24, 27-34]. Here, we ask whether linguistic diversity shows similar evolutionary dynamics. As diversification rates are known to vary within large language families [13-15], we calculate diversification rates between successive time periods,

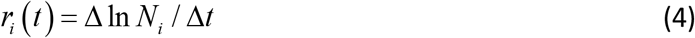

Therefore, for each phylogeny we calculate multiple growth rates and ask how they change with time, and in response to the number of languages. Because the time depth, *T*_*i*_, varies across the lineages between 1,000 to 13,000 years, and the number of languages within lineages varies from 5 to 1,268, we standardize time and size, so *N*_*i*_ (*t*)′ = *N*_*i*_ (*t*) / *N*_*i*,max_ and *t*_*i*_ ′ = *t*_*i*_ / *T*_*i*_. Therefore, *N*_*i*_ (*t*)′ is the proportion of the total number of languages at time, *t*, and time, *t*_*i*_ ′, is now relative to the time depth (or root age) of the phylogenetic tree (Figure S1). To calculate the rate of diversification of the trees we then divided the normalized sizes of the lineages in to ten bins of 0.1*t* ′ each, thus standardizing the number of measureable growth rates.

## Results

### Lineages through time

Lineages-through-time plots visualize the growth of lineages in a phylogeny [17]. Figure 2 shows the 9 phylogenies in our dataset that have topologies (colored lines) and the sum of all lineages across the 9 phylogenies (black line). Figure 2 shows that the expansion rate for the sum of all lineages *r* = 0.001 yr^-1^ (regression: *F*_1,064_ = 268,185, *r* ^2^ = 99.6%, *p* < 0.001). However, Figure 2 also shows that diversification rates vary both within and across language families. Some rates slow with time, others increase, while others remain stable. For example, diversification rates seem to decrease with time within the Austronesian and Bantu language lineages (white and red lines, respectively). Diversification rates in the Lezgic and Japonic lineages (pink and grey lines, respectively) seem to increase through time, and while the sum of all lineages combined (black line) seems to be relatively constant, there seems to be a slight decrease through time. Table 1 presents the expansion rates and doubling times for all lineages.

### Sum of all lineage diversification rates

To examine how overall diversification rates change through time we estimate the change in the sum of all lineages per millennia by averaging the data over 1,000 year bins and calculating the change over time. While the black line in Figure 2 is relatively linear on a semi-log scale, Figure 3A shows that the diversification rate changes nonlinearly over time as it is well-fit by a quadratic function (regression: *F*_8_ = 43.39, *r* ^2^ = 95%, *p* < 0.001). Diversity initially increases (i.e., *b* > *d*) until an inflection ∼5,000 BP after which diversity decreases toward the present (i.e., *b* < *d*). Figure 3B shows that the diversification rates are highly nonlinear with the number of languages, as shown by a quadratic function fit to the logged data, therefore capturing the skew (regression: *F*_8_ = 22.69, *r* ^2^ = 90%, *p* < 0.001): Diversification rates initially exhibit very strong positive density-dependence, where diversification rates increase rapidly with low number of languages, reaching maximum rates at 12 languages, thereafter declining steadily with increasing languages, i.e., negative density-dependence.

**Figure 3.**
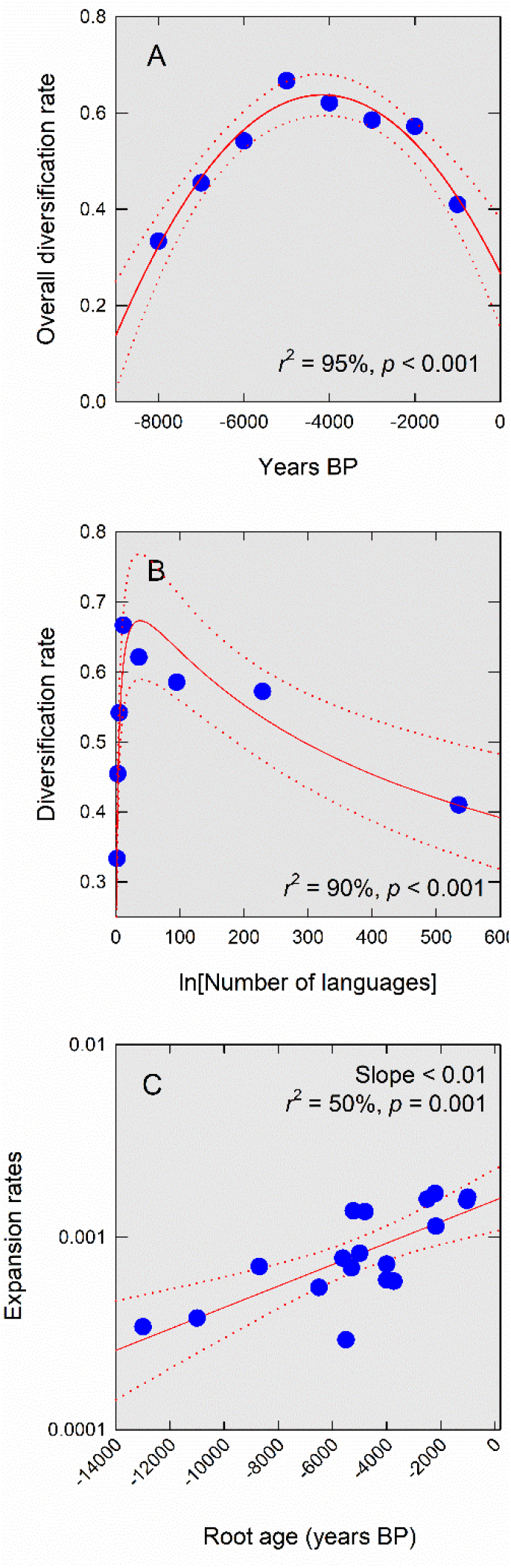
Diversification and expansion rates within and across lineages. A) Overall diversification rates over time fit with a quadratic function (red dotted lines are 95% confidence intervals around the slope). B) Overall diversification rates and the total number of languages, with a quadratic function fit to logged data. C) Expansion rates across lineages by their root ages fit by OLS regression.

### Expansion rates across lineages

Table 1 presents the expansion rates for all 18 lineages for which we have data. The mean rate of diversification across the full sample is 0.0001 yr^-1^ (bootstrapped 95% confidence interval = 0.00004 – 0.0001), or a doubling time of ∼770 years, similar to the estimated rate from the summed lineages. Figure 3C shows that the expansion rates of phylogenetic trees increases exponentially through time (regression: *F*_18_ = 28.84, *r* ^2^ = 54%, *p* < 0.001) by ∼20% per millennia, indicating that linguistic diversification is not constant through time.

Linguistic diversification rates across families appear to increase through time by an order of magnitude (Figure 3C). Taken at face value, diversification rates show an exponential increase over time with a slope of 0.0001. As such, the doubling time of diversification rates is about 7,000 years. However, this time dependency must be, at least in part, due to an unavoidable “pull of the present” effect [17,18]. This is because language families that arose recently have had less time to undergo extinction than older families. Moreover, our sample does not include many of the world’s smaller families and isolates. We can assume that these slow-growing languages would populate the lower right of Figure 3C to form a wedge shape to the graph and flatten the slope. For these reasons, an overall average diversification rate, as calculated in the previous paragraph, is a good estimate of the rate at which agricultural language families replaced hunter-gatherer language families around the world over the Holocene.

### Diversification rates within lineages

To examine the nature of diversification rates within lineages we recognize three basic forms of potential density-dependence; 1) exponential growth (i.e., density independence), slope = 0; 2) negative density-dependence, slope < 0; and 3) positive density-dependence, slope > 0. Figure 4 shows the same diversification rates for the nine lineages plotted as a function of the changing number of languages within lineages (statistics of each are given in the title of each panel). Four of the nine lineages show evidence of negative density-dependence (i.e., slopes > 0): these are the Austronesian, Bantu, Turkic, and Chapacuran phylogenies. The Lezgic lineage indicate positive density-dependence (slopes > 0). The Semitic, Uralic, Indo-European, and Japonic phylogenies exhibit exponential growth independent of density (slope = 0). Figures S2 and S3 in the online Supplementary Material show similar results by time.

**Figure 4.**
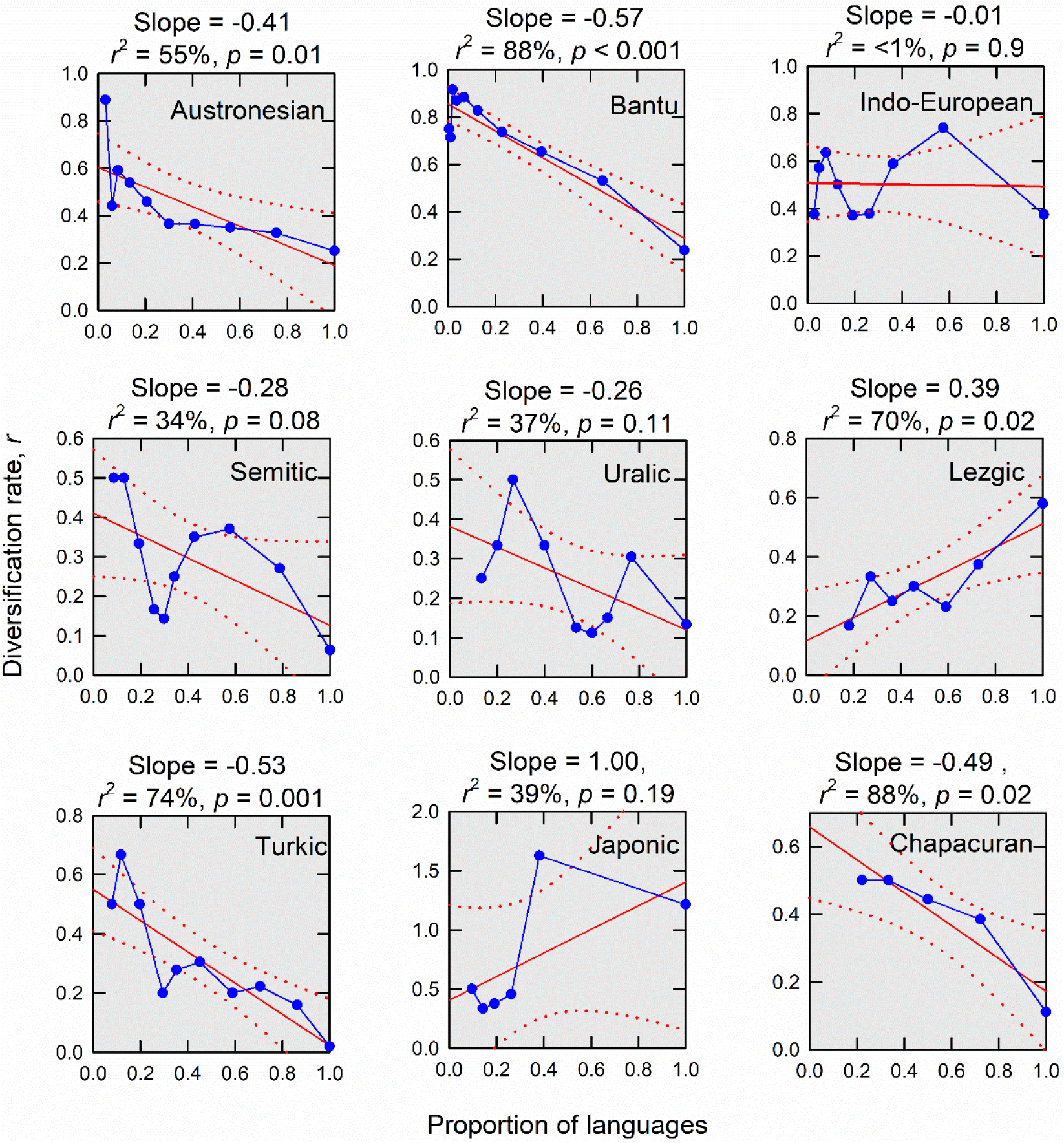
Different forms of density-dependent diversification rates among lineages. The Austronesian, Bantu, Turkic, and Chapacuran diversification rates are negatively density-dependent. Semitic, Uralic, Indo-European, and Japonic rates are not significantly different from constant, and therefore reflect exponential growth rates independent of density. Lezgic rates are positively density-dependent.

## Discussion

Our results indicate that rates of linguistic diversification are not constant through time, neither as a whole, nor within most of the individual lineages. Overall rates of diversification across all lineages initially increase rapidly for the first few thousand years of the Holocene and thus exhibit positive density-dependence where the actual rate of diversification increases with each new language speciated; diversity begets diversity. After the inflection point ∼5,000 BP, the diversification rate decreases, thus exhibiting negative density-dependence, likely reflecting increased competition and saturation of populations and landscapes as agricultural languages replaced hunter-gatherer languages. This highly nonlinear pattern describes an initial explosive burst of diversification as agricultural languages, and language families, spread rapidly throughout large parts of the planet over the course of the Holocene. Our results show strong evidence for similar patterns within some of the individua lineages where diversification rates are most rapid during the earliest stages of diversification (and at the smallest number of languages), indicating that as new lineages are born, they diversify explosively in punctuated bursts. Lezgic exhibits positive density-dependence, where, similar to the initial stages of overall diversity in the early Holocene, the rate of diversification increases through time toward the present and continues to diversify (i.e., diversity begets diversity). The slopes of the Indo-European, Semitic Uralic, and Japonic phylogenies are not significantly different from 0 and show no evidence of density-dependent diversification, suggesting that diversification rates have been approximately constant for 8,700 years.

The average expansion rate *r*_*E*_ of 0.001 yr^-1^ over all lineages means that the doubling time of the world’s linguistic diversity was on the order of ∼700 years as agricultural languages rapidly replaced hunter-gatherer languages. This diversification is driven by the underlying exponential growth rate of agricultural populations as they expanded from local geographic origins of domestication, eventually replacing most of the world’s hunter-gatherer populations [5-7]. When measured ethnographically, growth rates of natural-fertility human populations are often in the range of *r* ∼ 2 – 4% yr^-1^ [46,47]. Human populations therefore have the capacity to grow 20-40 times faster than languages diversify. However, the long-term growth rate of global human populations over the last 6,000 years, up until the industrial revolution, is also on the order of 0.001 yr^-1^ [48-50]. This suggests that language diversification and population growth rate on a global level are closely coupled and at near equilibrium levels (i.e., growth rates are only slightly above zero).

It might be the case that competition for limited resources forces local populations to diverge ethnolinguistically, resulting in some positive density dependence. Like species, linguistic change experiences punctuational bursts as languages go through splitting events [51,52]. Rates of new word gain have also been shown to be faster in larger populations [53], although others have argued for faster linguistic evolution in smaller populations [54]. A combination of larger populations and more competition could therefore fuel continuing ethnolinguistic diversification, but it remains an open question as to whether competition alone is sufficient to account for this dynamic. Islands are ideal test cases to establish whether ethnolinguistic diversity continues to increase once equilibrium population sizes have been reached. Indeed, linguistic diversity on islands has been shown to increase with time since initial settlement [55].

The rate at which agricultural populations expanded and replaced most of the world’s hunter-gatherer languages occurred at nonlinear, density dependence rates within most language families, where data are available. Like biodiversity in general, ethnolinguistic diversity may have continued to expand were it not for recent extinction events associated with globalization [56-59]. Interestingly, the data we present here suggests that the current language extinction crisis is part of a larger process of linguistic replacement that began in the mid-Holocene, ∼5,000 years ago with the expansion of agricultural language families, and now is driven by the expansion of global languages central to the expansion of global trade networks.

## Material and Methods

Linguistic phylogenies. We searched for the keywords “Bayesian linguistic phylogeny” to find studies that use Bayesian phylogenetic methods to attach known calendar dates to nodes of language trees so as to estimate root ages. We settled on 18 such linguistic phylogenies (Table 1). Of these we were able to obtain 9 of the actual consensus tree files with the highest posterior probabilities either from online supplementary material or by contacting corresponding authors. Unfortunately, we were not able to obtain error estimates for the consensus trees from the data made available.

Character-based phylogenetic methods using cognate codings of basic vocabulary words or phonetic data are useful for inferring the internal classifications and divergence dates of recent language family expansions. These methods are superior to traditional techniques in several ways by allowing for different rates of change between cognate sets and between different lineages, and by explicitly taking into account available archaeological dates and historical events to infer divergence dates [60]. While alternative phylogenetic methods exist [61-63], there is no competing alternative for our purposes because our analyses require a systematic approach that produces time-dated phylogenies.

The three largest language families in our sample are Austronesian [13], Bantu [14], and Indo-European [15]. Of these Indo-European has proven the most divisive with some arguing for the Kurgan expansion with homelands in the Pontic steppe dating 6,000 years ago versus others supporting the Anatolian expansion of farming around 8,700 years ago [7,15,64]. We opt for the latter following our inclusion criteria of Bayesian methods. Five families in our sample are Eurasiatic language families with estimated dates taken from Pagel and colleagues’ study of ultraconserved words [35]. These families are some of the oldest in our sample, and therefore generally considered the most controversial, but removing them from the sample yields only a slightly faster average diversification rate of 0.00104 yr^-1^. The remaining language families in the sample are more recent expansions (table 1), hence their divergence dates are likely more accurate.

Analyses. Lineage-through-time plots were created with the *ape* package in R. The sum of the lineages line in figure 2 is created by simply aligning the 9 available phylogenies together using absolute time as in the graph and adding all of the lineages together. How exactly each phylogeny relates to others at the basal node is not necessary information for the lineage-through-time plot since each additional lineage is simply counted at each point in time.

For within-family analyses of diversification rates as a function of time, we used the *R* package *TESS* [33]. This is a stochastic-branching model that flexibly fits time-varying speciation and extinction rates and accounts for incomplete sampling [34]. See Supplementary Material for complete details, *R* code, and tree data. The assumption of constant diversification rate *r* is a reasonable assumption for most of these language families so we made this assumption for each family in the full sample of 18. Diversification rates were estimated with equation 1 using the total (natural-logged) number of languages in a family from the Ethnologue [2], along with the estimated root age of the tree from the original studies (table 1). Doubling times were calculated as *ln*2/*r* [16]. While 4 of the phylogenies include dialects, we calculate diversification with counts of languages only.

## Author Contribution

Both authors, M.J.H. and R.S.W., contributed equally to this work.

